# Risk assessments underestimate threat of pesticides to wild bees

**DOI:** 10.1101/2023.09.15.557615

**Authors:** René S. Shahmohamadloo, Mathilde L. Tissier, Laura Melissa Guzman

**Author notes:** The authors contributed equally to this work. Corresponding author. (L.M.G.); (R.S.S.). **Author Contributions:** R.S.S., M.L.T., and L.M.G. designed the research, conducted the literature search, analyzed the data, and wrote the paper. **Competing Interest Statement:** The authors declare no competing interests.

## Abstract

Ecological risk assessments (ERA) are crucial when developing national strategies to manage adverse effects from pesticide exposure to natural populations. Yet, estimating risk with surrogate species in controlled laboratory studies jeopardizes the ERA process because natural populations exhibit intraspecific variation within and across species. Here, we investigate the extent to which the ERA process misestimates risk from pesticides on different species by conducting a meta-analysis of all records in the ECOTOX Knowledgebase for honey bees and wild bees exposed to neonicotinoids. We found the knowledgebase is largely populated by acute lethality data on the Western honey bee and exhibits within and across species variation in LD50 up to six orders of magnitude from neonicotinoid exposure. We challenge the reliability of surrogate species as predictors when extrapolating pesticide toxicity data to wild pollinators and recommend solutions to address the (a)biotic interactions occurring in nature that make such extrapolations unreliable in the ERA process.

**Synopsis:** Ecological risk assessments misestimate pesticide threats to pollinators sixfold by overextending acute lethality data on surrogate species to natural populations.

## 1. Introduction

The planetary scale of human impacts on nature has increased sharply since the 1970s, driven by the demands of a growing population that is challenged with mitigating and adapting to climate change (Díaz et al. 2019). Ecosystem deterioration and biodiversity loss are exacerbated by anthropogenic pollutants dispersing both on the land and in the water (Bongaarts 2019). One mechanism adopted by governments and regulatory agencies to mitigate biodiversity loss is the ecological risk assessment (ERA) process, which evaluates the likelihood and magnitude of risk that an environment might be impacted from exposure to one or more stressors (Levin et al. 1989). Reliably predicting how natural populations will respond to environmental stressors remains a defining feature of the ERA process (Levin et al. 1989).

Historically, the most influential metric of the ERA process has been the lethal dose at which a chemical will kill 50% of a test population (LD50) (Relyea & Hoverman 2006). This metric is derived from controlled laboratory studies quantifying toxicological responses on individual organisms (Barata et al. 1998; Stark et al. 2015). For over half a century, LD50s have guided regulatory agencies on how best to protect biodiversity while maintaining ecosystem services and economic output (USEPA 2014; Stark et al. 2015; Shahmohamadloo et al. 2022). Yet, environmental protection goals are designed to protect populations and communities, making it difficult to translate laboratory studies on individual organisms to interpretable field outcomes (Clements & Rohr 2009; Newman 2009; Hommen et al. 2010). This is especially true when assessing the effects of pesticides on target and nontarget species in agroecosystems, which must concomitantly suppress pests while having negligible and nontarget effects on beneficial species and humans (Stark et al. 2015).

Unsurprisingly, the ERA process for pesticides has come under scrutiny. Mounting criticisms (Chapman et al. 1998; Stark et al. 2004, 2015; Sánchez-Bayo 2014; Witwicka et al. 2022) challenge the preferential treatment of acute lethality data derived from stringent laboratory experiments on select surrogate species (EFSA 2013; Franklin & Raine 2019). Under the directives of the United States Environmental Protection Agency (USEPA) and similar regulatory agencies worldwide, toxicological responses of these surrogate species are used to extrapolate the ecological risks of pesticides using probabilistic approaches (USEPA 2004; NRC 2013; PMRA 2021). However, this process remains highly debated because surrogate species may not be reliable predictors of what may happen to thousands of beneficial species exposed to pesticides in nature (Banks et al. 2010, 2011; Franklin & Raine 2019; Siviter et al. 2021; Witwicka et al. 2022). Additional considerations, including associated effects on heterogeneous and genetically diverse populations (Forfert et al. 2017), interactions between pesticides and climate change (Delcour et al. 2015), and the metabolic effects of pesticides on species life-history traits (Cook 2019), to name but a few, add further complexities unaccounted for in the ERA process ostensibly designed to assess risks to natural populations. Paradoxically, the pursuit to assess comparative toxicity between species and understand the mechanisms of toxicity require maximal control of laboratory conditions. This approach ignores the complexity of interactions between pesticides and other (a)biotic stressors in nature, an issue that reduces confidence in extrapolating between lab and field studies and yields considerable uncertainty among regulators in pesticide risk management (Monchanin et al. 2019).

Here, we investigate the extent to which LD50 assays on surrogate species represent the distribution of LD50s for related species. We profile the most widely used insecticides in the world (Sánchez-Bayo 2014) and their documented toxicity on a pollinator driving policy reform: neonicotinoids and the honey bee (*Apis*). As ‘hidden killers’ applied to seeds and soils (Sánchez-Bayo 2014), neonicotinoids have gradually contributed to widespread collapse of domestic honey bee colonies over decades (Gross 2013). Demands across all sectors to protect bees —and more broadly pollinators as providers of vital ecosystem services— led to concerted efforts by governments around the world to utilize the Western honey bee (*Apis mellifera*) as a surrogate for both *Apis* and non-*Apis* bees (Thompson & Pamminger 2019) via a three-tiered approach embedded in the ERA process (*SI Appendix*, SI 1). We conducted a meta-analysis of all acute and chronic lethality records in the ECOTOX Knowledgebase (hereafter, called ECOTOX) for honey bees exposed to neonicotinoids, the world’s largest curated database of ecologically relevant toxicity tests to support environmental research and risk assessment (Olker et al. 2022). This knowledgebase is used by the USEPA and regulatory agencies worldwide in pollinator risk assessment, risk management, and research. We collated all LD50 data along with any accompanying environmental parameters measured in ECOTOX (e.g., origin of bee strain, min and max temperature, duration of studies, and route of neonicotinoid exposure, to name a few). We hypothesized that regulatory agencies are overextending lethality data of the Western honey bee to other *Apis* and non-*Apis* species in risk assessments and do not sufficiently represent the breadth of intraspecific variation under more realistic conditions.

## 2. Methods

All records from ECOTOX https://cfpub.epa.gov/ecotox/; database access October 16, 2022) were downloaded with filters applied for neonicotinoids and bees. We evaluated the variation in LD50 both within *Apis* and between *Apis* and non-*Apis* species, and subsetted assays that reported an LD50. The LD50s reported across all assays were standardized to ng/organism, which were used to generate species and genus sensitivity curves. This resulted in a total of 252 assays that evaluated LD50 on any bee species. We returned to each original study and recorded the sample size for each assay. We obtained sample sizes for 226 assays. Further, ECOTOX contains information on 95% confidence intervals. For some studies where the 95% confidence intervals were missing from ECOTOX, we were able to obtain from the original studies. In total, we had 162 assays with confidence intervals. For the remaining assays we were able to obtain the standard error of the mean response for 1 assay, standard deviation (which we then converted into standard error by standardizing the units and combining with the sample size) for 18 assays, and finally 29 assays had a chi-squared and *p*-value. In total, we obtained 210 assays with a mean response, sample size, and some measure of variation. For the studies with *p*-values, we constrained these *p*-values to be between 0.0001 and 0.9999 to allow for meta-analysis convergence.

We evaluated the inter-genus variation in LD50 via a mixed-effects meta-analysis regression using the *meta* package in R (Balduzzi et al. 2019), with genus and the duration of the study (in days) as predictors. All LD50s were log transformed to meet normality assumptions. Each pesticide and route of exposure was evaluated separately.

We then selected *A. mellifera* and imidacloprid assays —the most commonly tested across all assays— for a case study to determine whether records from ECOTOX accounted for genetic and environmental interactions with the toxicant. Relevant parameters investigated were: the origin of bee strain; route of neonicotinoid exposure to bees; and min and max temperature. For all analysis the *p*-level significance cutoff was 0.05. See *SI Appendix* (SI 2-5) for further details.

## 3. Results

### 3.1 The ECOTOX Knowledgebase

We identified 252 assays spanning 49 studies from ECOTOX to generate the LD50 species and genus sensitivity curves (Dataset S1). The bee genera included were *Apis* (200/252 assays; 38/49 studies), *Bombus* (24/252 assays; 8/49 studies), *Megachile* (7/252 assays; 1/49 studies), *Melipona* (3/252 assays; 2/49 studies), *Nannotrigona* (2/252 assays; 1/49 studies), *Osmia* (7/252 assays; 3/49 studies), *Partamona* (1/252 assays; 1/49 studies), *Plebeia* (1/252 assays; 1/49 studies), *Scaptotrigona* (4/252 assays; 2/49 studies), and *Tetragonisca* (3/252 assays; 2/49 studies). The route of neonicotinoid exposure also differed among studies in ECOTOX, which we broadly categorized via ‘dietary’ (diet = 8/252 assays; food = 162/252 assays) and under ‘topical’ (topical = 28/252 assays; dermal = 41/252 assays; environmental = 3/252 assays; spray = 9/252 assays).

For the case study on imidacloprid and *A. mellifera*, no studies investigated the role of genetic diversity on phenotypic outcomes. In addition, although reporting the origin of a colony is a requirement for studies in Europe on the risk assessment of plant protection products on bees (EFSA 2013), only 77/129 studies reported a location where their Western honey bee colony originated; yet, upon closer inspection, none of these studies reported strain-specific information. Regarding biotic and abiotic variables that could influence bee response to neonicotinoid exposure (Cook 2019), they were considered in a very small portion of the studies from ECOTOX. Less than 6% of the case studies considered diet as potentially affecting honey bee responses to imidacloprid, despite evidence of an interaction between topical and dietary exposure to neonicotinoids (Tosi et al. 2017; Leza et al. 2018; Stuligross & Williams 2020; Klaus et al. 2021; Costa et al. 2022). Nutritional and metabolic properties of neonicotinoids were not investigated in the studies while min and max temperatures were reported in solely 13/129 studies. Finally, only one study considered the impact of imidacloprid on reproductive output of the queen bee. Because so few studies evaluated the impact on reproduction or the role of genetic diversity, nutritional properties, metabolism, and temperature, we do not examine this data further.

### 3.2 LD50 variation within and across bees

To test whether LD50 varies due to genetic diversity within and across species, we ran a meta-analytical regression, and compared genus level estimates for all bee genera to those of *Apis*. Because the predictor is categorical (bee genera) the first group is treated as the intercept, which in this case was *Apis*. We did not conduct post-hoc comparisons because we were mainly interested in the comparison between all genera to *Apis* given it is a surrogate species in toxicity testing. The model also provides an estimate for differences between each genus and the intercept; a negative estimate, for instance, shows the LD50 of that genus is lower compared to *Apis*, while a positive estimate shows the LD50 of that genus is higher than *Apis*. A lower estimate therefore suggests that toxicity increases compared to *Apis*. We also evaluated the effect of the study duration using the number of days of the study. A negative estimate for duration indicates the LD50 across all groups decreases as duration of the study increases (i.e., toxicity significantly increases).

For dietary exposure, the model shows that bee genera exhibited significant variation in LD50 response to neonicotinoids, both within and across genera (Figure 1; Dataset S1). *Bombus* was significantly more sensitive to imidacloprid and thiamethoxam than *Apis* (estimate = -6.03, se = 1.73, *p*-value = 0.00050; estimate = -2.02, se = 0.80, *p*-value = 0.011). *Megachile* was significantly more sensitive to clothianidin, imidacloprid, and thiamethoxam than *Apis* (estimate = -7.37, se = 1.42, *p*-value < 0.0001; estimate = -4.20, se = 2.12, *p*-value = 0.048; estimate = - 4.99, se = 1.19, *p*-value < 0.0001). *Partamona* was significantly more sensitive to imidacloprid than *Apis* (estimate = -6.49, se = 2.71, *p*-value < 0.017). All estimates from dietary exposure to neonicotinoids were also negative, demonstrating toxicity significantly increases as the duration of studies increases. As for variation within species, we found *Apis* exhibits variation in LD50 up to six orders of magnitude across all neonicotinoids (Figure 1).

**Figure 1.**
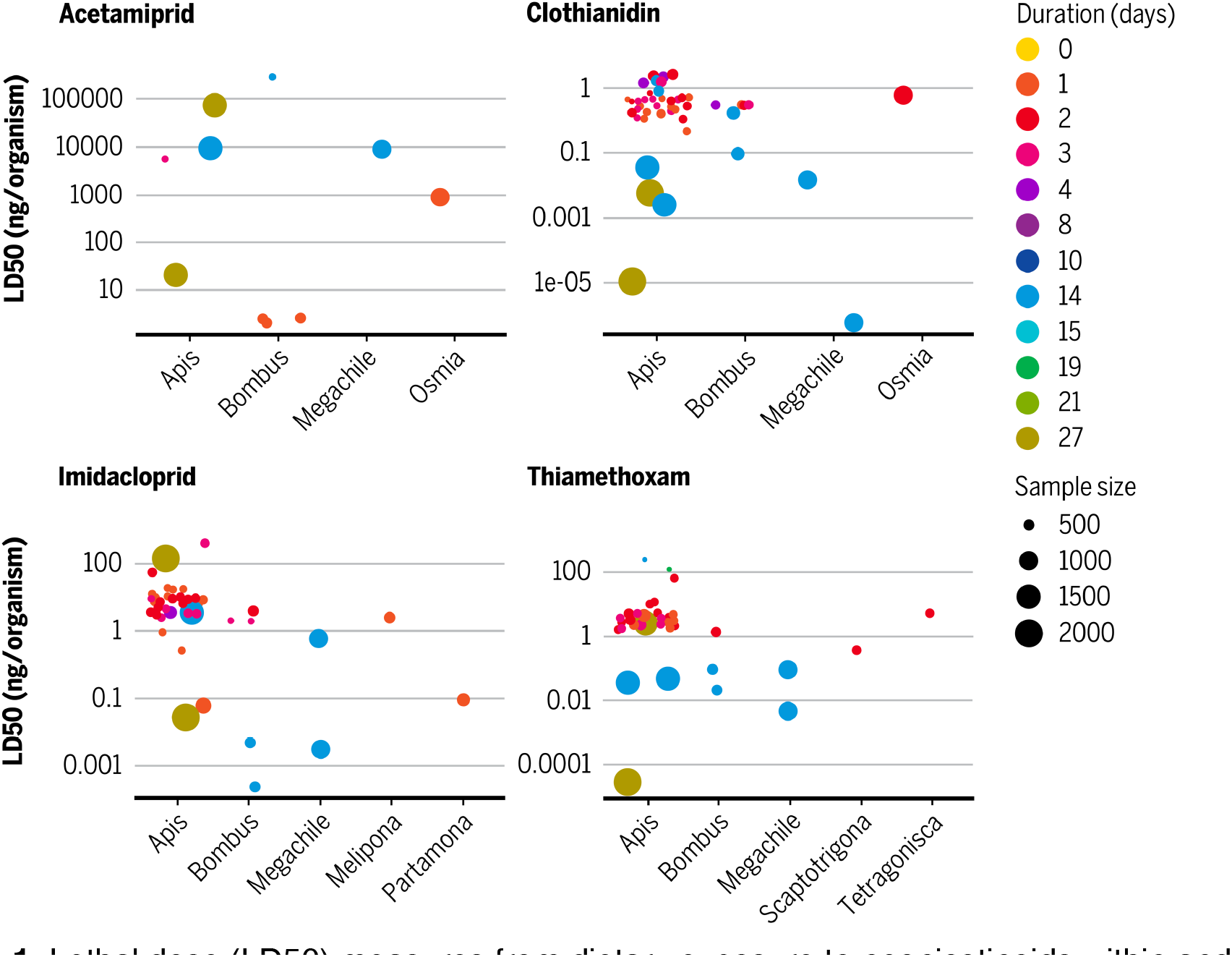
Lethal dose (LD50) measures from dietary exposure to neonicotinoids within and across bees. The surrogate, *Apis*, is compared to wild bees of seven genera (*Bombus, Megachile, Melipona, Osmia, Partamona, Scaptotrigona*, and *Tetragonisca*). Duration of studies are color-coated and sample size is indicated by the diameter of the circle.

For topical exposure, the model similarly shows that bee genera exhibited significant variation in LD50 response to neonicotinoids, both within and across genera (Figure 2; Dataset S1). Several genera were significantly more sensitive to imidacloprid than *Apis*, including *Melipona* (estimate = -3.09, se = 0.85, *p*-value = 0.00028), *Nannotrigona* (estimate = -3.43, se = 1.15, *p*-value = 0.0029), while *Osmia* was significantly less sensitive to imidacloprid (estimate = 4.46, se = 1.25, *p*-value = 0.00035). Estimates from topical exposure to neonicotinoids were also negative, demonstrating toxicity significantly increases as the duration of studies increases. As for variation within species, we found *Apis* exhibits variation in LD50 up to four orders of magnitude across all neonicotinoids (Figure 2).

**Figure 2.**
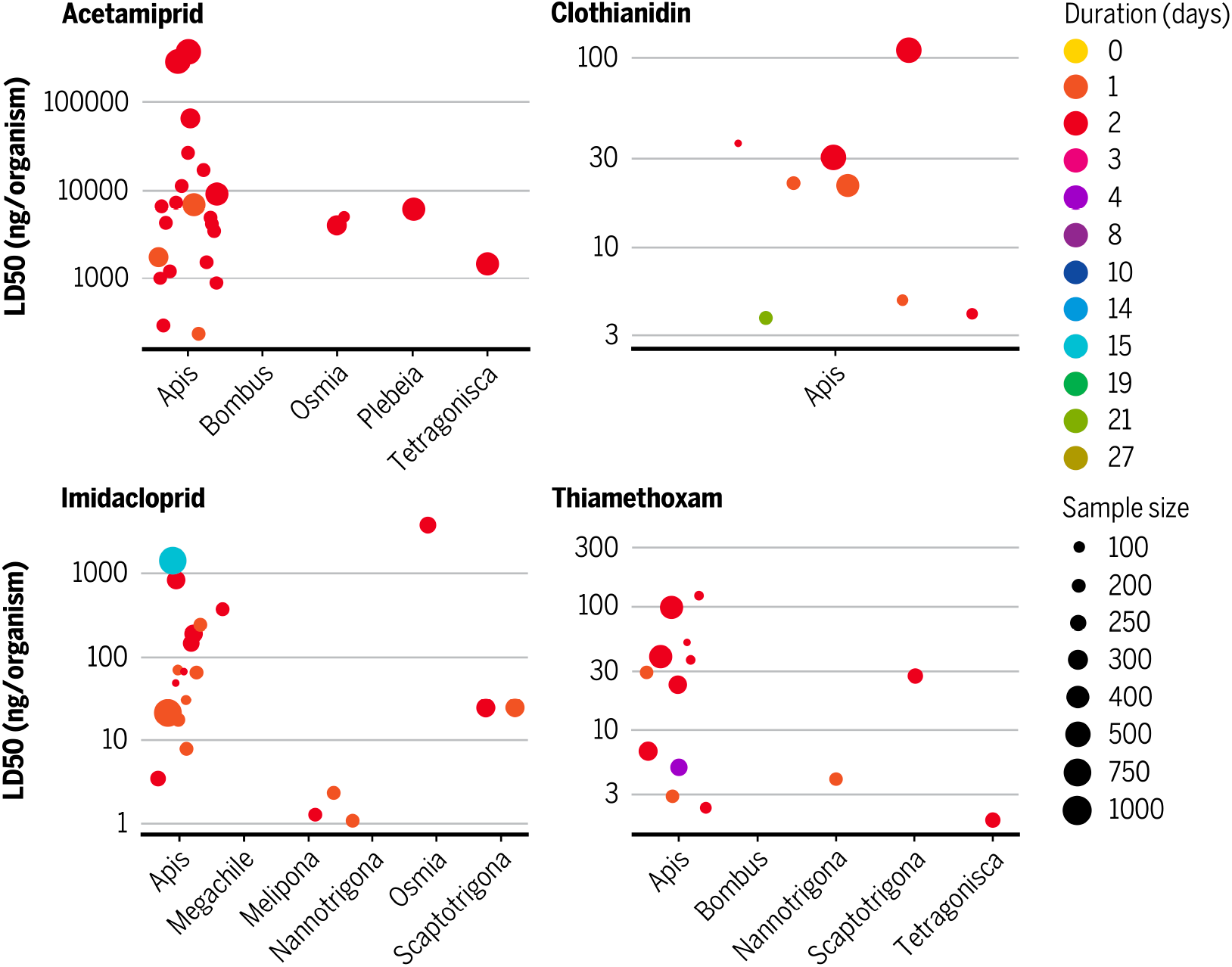
Lethal dose (LD50) measures from topical exposure to neonicotinoids within and across bees. The surrogate, *Apis*, is compared to wild bees of eight genera (*Bombus, Megachile, Melipona, Nannotrigona, Osmia, Plebeia, Scaptotrigona*, and *Tetragonisca*). Duration of studies are color-coated and sample size is indicated by the diameter of the circle.

### 3.3 Duration of studies

Aware that the ECOTOX Knowledgebase is largely populated by acute lethality data between 1-5 days —particularly topic studies (Figure 2)— we further tested whether LD50 is influenced by the duration of studies. Imidacloprid LD50s from topical exposure were significantly influenced by the duration of studies (estimate = 0.26, se = 0.11, *p*-value <0.019). The estimate from imidacloprid topically exposed was also positive. This means that toxicity decreases with the duration of studies, and becomes less lethal more rapidly. Although for imidacloprid only — owing to the fact that most studies in ECOTOX conducted assays with this neonicotinoid— this result nonetheless illuminates the importance of chronic versus acute studies on pesticide toxicity in bees.

## 4. Discussion

### 4.1 The impact of interspecific variation on pollinator risk assessment

Pollinator risk assessments are predicated on studies that provide key lethality data for regulators to make decisions invested in protecting biodiversity while maintaining ecosystem services and economic output (Levin et al. 1989; USEPA 2004; Relyea & Hoverman 2006; Stark et al. 2015; Bongaarts 2019; Shahmohamadloo et al. 2022). The three-tiered approach adopted by the USEPA and other regulatory agencies worldwide provides a framework by which regulators translate study results to inform their decisions on pesticide risk management (Olker et al. 2022). For neonicotinoids and bees, the majority of studies found in the ECOTOX Knowledgebase used *Apis* and most were conducted in laboratories on an acute timescale. Although *Apis* is a model organism for pollinator risk assessment, growing concerns advocate it is giving false confidence on pesticide safety thresholds for other bee species as life histories and route of exposure across genera may vary widely (Boyle et al. 2019; Franklin & Raine 2019; Schmolke et al. 2021; Witwicka et al. 2022).

We validate these concerns by showing *Apis* does not accurately estimate lethality risks from dietary (Figure 1) or topical (Figure 2) exposure to neonicotinoids on native bees from the genera *Bombus, Megachile, Melipona, Nannotrigona, Partamona*, and *Osmia. Apis* makes up the majority of assays (79.4%) in ECOTOX, while native bees lag behind (20.6%). Trait-based vulnerability analyses suggest non-*Apis* bees may be more vulnerable to pesticides than the Western honey bee due to traits impacting exposure and population recovery potential (Schmolke et al. 2021). Recent findings showing differences in expression of the neonicotinoid target receptor nicotinic acetylcholine receptor (nAChR) among tissues and life stages within species (*A. mellifera*) and between species (*A. mellifera* and *Bombus terrestris*) bolster our concerns about generalizing toxicity estimates across bees and highlights the need for a greater understanding of the molecular targets under selection and variation in those targets (Witwicka et al. 2022). As one example, Europe is moving towards a holistic approach to risk assessment that acknowledges non-*Apis* species as integral to pollination services and wild bee diversity (More et al. 2021; Knapp et al. 2023). For North America and other regions, diligent work is needed to develop toxicity assays for native and wild bees who, as we have found, differ in their life histories and sensitivities to neonicotinoids by several orders of magnitude compared to the Western honey bee.

Native bee sensitivities in LD50 responses to neonicotinoids are questionable when the duration of studies increases from acute (<5 days) to chronic (>10 days). Demands for risk assessments to include more chronic toxicity studies are ongoing and widely debated for pesticides (Sanchez-Bayo & Goka 2014; Sánchez-Bayo & Tennekes 2017; Tsvetkov et al. 2017; Stuligross & Williams 2021). Arguments in favor of maintaining status quo suggest ‘Tier I’ studies in honey bee acute risk assessment guidelines are protective of other bee species and tests should only be conducted in cases of concern (Thompson & Pamminger 2019) — a conclusion drawn from studies not found in ECOTOX. As is the case from our meta-analysis, parsing the route of toxicity uncovered a large bias of acute assays for topical studies. The chronic impacts on bees exposed topically remain largely unknown, and more chronic studies are needed for dietary exposure. Our findings should compel risk assessors to question whether the LD50 responses measured in *Apis* on an acute timescale meets the broader objectives of protecting wild bees and biodiversity chronically exposed to pesticides.

### 4.2 Sources of intraspecific variation within Apis

The wide variation in LD50 up to six orders of magnitude within *Apis* (Figure 1, 2) brings into question whether lethal doses are truly knowable. Several factors likely contributed to this variation. Below we highlight three factors —namely genetic variation, temperature, and nutrition— which we believe are synergistic and deserve special consideration in the ERA process, and more broadly biodiversity conservation efforts.

Failure to incorporate intraspecific genetic variation in toxicity studies hinders the estimation and understanding of genetic diversity, as well as the representativeness of the tested strain within a species or genus (Shahmohamadloo et al. 2023). This limitation introduces significant uncertainty when applying laboratory results to real-world scenarios, which undermines the accuracy and effectiveness of risk management strategies (Forbes 1998; Coutellec & Barata 2011; Thompson et al. 2023). Oversight of intraspecific variation becomes particularly relevant in instances of pesticide resistance evolution, as the presence of genetic variation can significantly reduce the effectiveness of agrochemicals due to natural selection (Hawkins et al. 2019). Thus, evolutionary processes such as adaptation to pesticides can plausibly affect the outcome of toxicity tests (Coutellec & Barata 2011) and need better representation in risk assessments. If the ERA process cannot quantify interactions between toxicants and evolutionary forces on a population scale (e.g., genetic drift, inbreeding, selection), an accompanying decrease in genetic diversity may have negative effects at the population level, including: a reduced ability of populations to evolve genetically-based changes in traits in response to environmental fluctuations (Thompson et al. 2023); increased mutations (Coutellec & Barata 2011); risks of extinction (Exposito-Alonso et al. 2022); and pesticide resistance (Gould et al. 2018). Pesticides act as a strong agent of natural selection, often capable of driving rapid adaptation (Gould et al. 2018; Rudman et al. 2018). The standing stock of intraspecific genetic variation as it sits today can be important in predicting pesticide responses because there are differences in genotype. Yet it is reasonably difficult to predict evolutionary responses in nature where the selective landscape has multiple drivers of natural selection causing phenotypic variation, an outward expression of the genotype (Endler 1986). Lack of strain-specific, genetic information from the studies we reviewed in ECOTOX represent a major gap and further demonstrates the need for toxicity testing to assess the additive genetic variance in response to pesticides. Additive genetic variance is measurable at a population level and, when combined with information about natural selection, can be used to predict the pace and direction of adaptation (Rudman et al. 2022). Collecting this data represents a crucial responsibility for scientists and more broadly supporting the ERA process.

Although the ERA process in its current state does not explicitly incorporate the interactive effects of climate and pesticides, there is growing recognition that temperature can affect the risks of pesticides to pollinators. For instance, honey bees were more sensitive to imidacloprid and thiamethoxam at 24°C than 35°C (Saleem et al. 2020). This reduction in toxicity at 35°C was not due to degradation of the pesticide itself; nonlinearities may arise when heat stress is high enough to induce physiological responses that allow bees to better tolerate neonicotinoids (2020). Increases in temperature can also affect the absorption and metabolism of pesticides. For instance, studies on the effect of organophosphorus insecticides on midge (*Chironomus tentans*) lethality showed that acute toxicity increased with temperature, likely due to the metabolic activation of organophosphorus insecticides to more toxic forms (Lydy et al. 1999). However, the trend may not always be straightforward. Studies evaluating the combined effect of thiamethoxam and temperature on homing performance of honey bees found that lower temperatures (16–20°C) exacerbated the homing failure induced by the pesticide (Monchanin et al. 2019). Further, the interaction between temperature and neonicotinoids may depend on the type of behavior being observed. Neonicotinoids had a greater effect on movement and food consumption at a low temperature (21°C) yet a greater effect on flight distance at a high temperature (30°C) (Kenna et al. 2023). Considering 13/129 studies reported temperature from our analysis of ECOTOX, monitoring temperature can be critical to interpreting data used for risk assessments.

In their natural environment, bees are primarily exposed to neonicotinoids through the diet (e.g., nectar, pollen, water consumption), although ground-nesting bees can also be exposed to neonicotinoids and their associated byproducts that leach into soils (Main et al. 2020; Willis Chan & Raine 2021). Dietary exposure to neonicotinoids comprised 70% of studies in ECOTOX for honey bees. Although most studies assessed honey bee lethality or behavioral responses following dietary exposure to imidacloprid specifically, they overlooked the interactions of these molecules with the diet, its nutrient content or the presence of other biomolecules. This could help explain the observed differences between topical and dietary exposure (Figure 1, 2). This observation is of special concern given nutritional stress substantially increases bee sensitivity to pesticides, including neonicotinoids (Tosi et al. 2017; Leza et al. 2018; Stuligross & Williams 2020), while an appropriate diet can alleviate or even buffer the detrimental effects of insecticides on bees (Klaus et al. 2021; Costa et al. 2022). Interestingly, primary or secondary compounds of plants (e.g., alkaloids or flavonoids) can help bee detoxification processes (Johnson et al. 2012; Balieira et al. 2018), ultimately offsetting negative impacts of insecticides on bee physiology, health and performance (Costa et al. 2022; Riveros & Gronenberg 2022). Thus, controlled studies looking at bee responses to a pesticide in a nutritionally-rich setting may not capture the complexity of pesticide exposure in the natural environment.

## 5. Recommendations

Our meta-analysis of the ECOTOX Knowledgebase uncovers substantial areas of development in the interpretation of pesticide risks to pollinators based on data largely derived from the surrogate species, *Apis*. Below we offer three recommendations, in writing and illustration (Figure 3), which we anticipate will help scientists, regulatory agencies, and policy-makers in achieving their aims to protect biodiversity and ecosystem services, while maintaining economic outputs.

**Figure 3.**
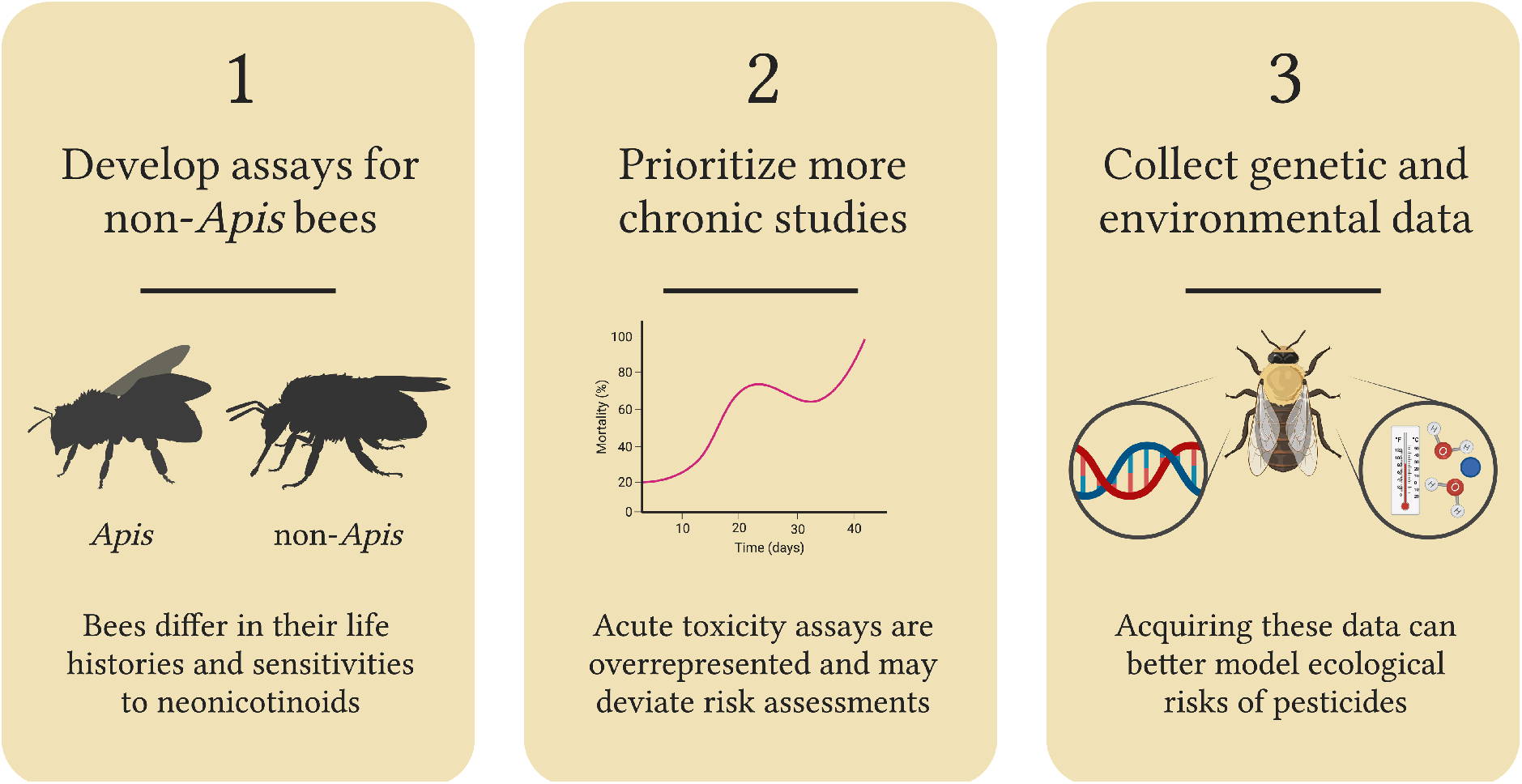
Recommendations to scientists, regulatory agencies, and policy-makers overseeing ecological risk assessments of neonicotinoids on bees.

- **Develop assays for non-*Apis* bees**. The ERA process relies heavily on *Apis* to predict risks from pesticides to other bees. Yet, our meta-analysis shows North American native bees differ in their life histories and sensitivities to neonicotinoids by several orders of magnitude compared to *Apis*. Developing toxicity assays for wild native bees and integrating these into the ERA process is a promising step forward that acknowledges non-*Apis* species as integral to pollination services and wild bee diversity.
- **Prioritize more chronic studies**. The ERA process for pollinators exposed to neonicotinoids relies heavily on acute toxicity data. Our meta-analysis brings into question whether LD50 responses measured on an acute timescale meet the broader objectives of protecting native bees and biodiversity chronically exposed to pesticides. Chronic studies should be prioritized.
- **Collect genetic and environmental data**. Studies need to better account for and report genetic and environmental data (e.g., climate and nutrition), otherwise it becomes difficult to interpret LD50s. Acquiring this data can assist with developing better models to characterize ecological risks and developing better mitigation strategies.

## Acknowledgments

This work was supported by Liber Ero Postdoctoral Fellowships, an NSERC Postdoctoral Fellowship, and the USC Wrigley Faculty Innovation Award. We thank S. Otto, W. Palen, P. Sibley, S. Rudman, and K. Cox for their constructive comments and discussions.

