## Supporting Information for "Risk assessments underestimate threat of pesticides to wild bees"

**Number of pages: 5**

Table of Contents | Page

| Section | Page |
|---|---|
| SI 1. Ecological risk assessment guidelines for neonicotinoids | S3-4 |
| SI 2. Data access | S4 |
| SI 3. Species and genus sensitivity curves | S4 |
| SI 4. Case study on imidacloprid and Apis mellifera | S5 |
| SI 5. Statistical analysis | S5 |
| SI 6. Dataset S1 | S5 |
| Supporting References | S5 |

***SI 1. Ecological risk assessment guidelines for neonicotinoids***

When the USEPA and PMRA conduct an ecological risk assessment of a pesticide to pollinators, it considers both the level of risk and the feasibility of mitigation measures needed to reduce that risk. If the risk is deemed high, regulatory agencies may require that specific mitigation measures be put in place to reduce the risk to an acceptable level. These mitigation measures could include changing the timing of spraying, using specific application methods, or requiring buffer zones around sensitive areas. The goal is to find a balance between protecting pollinators and ensuring that farmers and other users can continue using pesticides to protect their crops and livelihoods.

At the core of the risk assessment guidelines is a three-tiered approach that estimates toxicity from environmental concentrations derived from laboratory (Tier I), semi-field (Tier II) or field (Tier III) studies. All three tiers are meant to generate conservative estimates of risk and reduce uncertainties related to pesticide risks.

**Tier I studies:** Laboratory acute and chronic toxicity studies with adult and larval honey bees exposed by oral and contact routes, to derive an LD50 metric.

**Tier II studies:** Semi-field tunnel and feeding toxicity studies where honey bee hives are exposed to neonicotinoids in enclosed tunnels with treated crops, or where whole colonies are tested for feeding in a laboratory. Residue studies can also be employed to assess neonicotinoid levels on pollen and nectar from crop application.

**Tier III studies:** Field studies that are intended to evaluate the full potential effects on colonies under actual use conditions. These studies are designed to assess risk hypotheses or uncertainties. The goal of these studies is to reduce the confounding effects, and therefore the duration and site locations are carefully considered. If a study aims to evaluate an overwintering component, then a longer study duration is important. However, longer duration in the study could cause an increase in natural variation among replicates, which can obscure the effects of pesticides.

Following the three-tiered approach, risk quotients are calculated for individuals bees following these steps:

**Step 1. Determine if bees are exposed.** In the case of pesticides, the first step is assessing whether bees could have been exposed. Outdoor spray applications are assumed to result in exposure for adult bees. Other life stages can be exposed while foraging. Exposure is also assessed depending on when the bees are likely to be foraging.

**Step 2. Calculate Tier I Screening-Level Risks.** If bees are expected to be exposed, then estimated exposure concentrations (EECs) are compared with Tier I acute and chronic levels of toxicity using individual bees exposed in the laboratory (e.g., LD50). Specifically, risk quotients (RQ) depend on the model that generates the EECs.

**Step 3. Refine Tier I Screening-Level Risk Estimates.** If risk concerns are higher, then the risk assessment made in step 2 can be refined with additional data. These refinements can include residue data available from crops or pesticide residues in pollen and nectar.

**Step 4. Consider Uncertainties, Risk Mitigation Options and Need for Tier II Risk Estimation.** If even after refinement, risks are still identified, then mitigation options might be considered such as reduction in application rates (which would modify the exposure concentrations) or restriction of the application methods. Modifications of the timing of pesticide application can also minimize the exposure to bees. Tier II studies may also be warranted either to identify and quantify exposure specifically on pollen/nectar or evaluate pesticide effects at the whole colony level under semi-field conditions. Tier II studies therefore reduce uncertainty associated from extrapolating results on individual bees under laboratory conditions.

**Step 5. Consider Uncertainties, Risk Mitigation Options and Need for Tier III Studies.** Based on the risks identified under Tiers I and II, risk mitigation options are considered such as reduced application rates, reduced application intervals, restrictions during blooming time or off-labeling use. Further studies can also be conducted to improve uncertainties. Tier III studies are full-field studies designed to mimic actual pesticide applications and exposures in the environment.

### *SI 2. Data access*

All records from the ECOTOX Knowledgebase (1) (https://cfpub.epa.gov/ecotox/; knowledgebase access October 16, 2022) were downloaded with the following filters applied: Realm: terrestrial; Chemicals: Neonicotinoids, including Acetamiprid - 135410207, Clothianidin - 210880925, Dinotefuran - 165252700, Imidacloprid - 138261413, Imidaclothiz - 105843365, Nitenpyram - 150824478, Nithiazine - 58842209, Paichongding - 948994169, Thiacloprid - 111988499, and Thiamethoxam - 153719234; Species: Insects/Spiders. These filters resulted in 13,762 assays obtained from 1,048 studies.

### *SI 3. Species and genus sensitivity curves*

We evaluated the variation in LD50 both within *Apis* and between *Apis* and non-*Apis* species. We 1) subsetted the 13,762 assays to those that evaluated any bee species, and 2) further subsetted results that reported an LD50. For each species, we found the family name using *taxize* (2) and filtered observations for the following families: *Apidae*, *Megachilidae*, and *Halictidae.* No studies found in the ECOTOX Knowledgebase have been performed on *Melittidae*, *Andrenidae*, *Stenotritidae*, or *Colletidae.* In summary, from the 13,762 assays 4,734 were performed on any bee species (including *Apis* and non-*Apis*). From the 4,734 assays, 351 of those evaluated the LD50. LD50s reported across all assays were standardized to ng/organism due to wide variability in units that measured toxicity. We standardized the following units to ng/org: AI ng/org, AI ug/org, mg/bee, ng/org, pg/org, ug/bee, ug/org, ul/org, consisting of 279 assays. The following units were unable to be standardized: AI lb/acre, AI ppm, AI ppm diet, AI ug/org/d, mg/L, ng/ul, ug/ml, consisting of 72 assays. Of those 72 assays, 16 were replicate results reported in different units). After standardization, we derived endpoints for LD50 (279 assays). The LD50 values were distributed among the following neonicotinoids: acetamiprid (42 assays), clothianidin (64 assays), imidacloprid (73 assays), and thiamethoxam (73 assays). The remaining neonicotinoids could not be included in the data analysis due to insufficient LD50 values from the ECOTOX Knowledgebase (dinotefuran, 15 assays; nitenpyram, 1 assay; thiacloprid, 11 assays). This resulted in a total of 252 assays that evaluated LD50 on any bee species. Studies that reported min, max, and mean observed responses were used to compare the range of toxin sensitivity across bee species.

### *SI 4. Case study on imidacloprid and Apis mellifera*

To evaluate whether records from the ECOTOX Knowledgebase accounted for genetic and environmental interactions with the toxicant, we examined all laboratory and field studies that tested the effect of imidacloprid on *Apis mellifera*. This neonicotinoid and species were chosen because they were the most commonly tested (1,377 of 13,762 assays).

The 1,377 assays spanned 129 studies; 108 were in the laboratory, 15 were in natural field conditions, and six were in artificial field conditions. From the 108 laboratory studies, only 102 were analyzable (five not found or accessible online and one not peer-reviewed); for the 21 natural and artificial field studies, only 18 were analyzable (two not found online and one proceedings of a conference inaccessible). Additional parameters including the origin of bee strain, route of neonicotinoid exposure to bees, min and max temperature, humidity, study location, and metabolic properties of neonicotinoids were collated from all publications. Although none of these studies tested the effect of genetic variation on the response variable (e.g., LD50), by investigating the source of the bee strain we were able to determine where these bees originated and how much genetic variation was tested.

### *SI 5. Statistical analysis*

We evaluated the inter-genus variation in LD50 via a mixed-effects meta-analysis regression, with genus and the duration of the study (in days) as predictors. All LD50s were log transformed to meet normality assumptions. Each pesticide was evaluated separately. Out of the 252 assays, only 113 had reported CI in the knowledgebase. For each study, we collated the sample size from the original papers and where possible acquired either the standard deviation or standard error. This was only possible for a handful of studies as the majority did not report any measure of variation, or reported it in the form of graphs but not in the actual results. Corresponding authors from these studies were additionally contacted and given 30 days to provide the requested information above. Therefore, to analyze these studies in a standard way, we inferred standard error from the sample size alone. We used the *meta* package (3).

For all analysis the *p*-level significance cut-off was 0.05. All analyses were completed in R version 4.2.2 (4) and can be retrieved here: https://github.com/lmguzman/bee_pesticide_epa

### *SI 6. Dataset S1*

A spreadsheet detailing all data retrieved from the ECOTOX Knowledgebase is provided here (sheet named ‘Dataset S1’).
